# Human genetic variation shapes the response of neurons to interferons

**DOI:** 10.1101/2025.05.28.653507

**Authors:** Milena M Andzelm, Sonia Bolshakova, Noah Pettinari, Matthew Tegtmeyer, Dan Meyer, Autumn Johnson, Curtis J Mello, Connie T Yu, Ilinca Mazureac, Giulio Genovese, Adrianna Maglieri, Kiku Ichihara, Marina Hogan, Derek Hawes, Steven A McCarroll, Ralda Nehme

**Affiliations:** Stanley Center for Psychiatric Research, Broad Institute of MIT and Harvard, Cambridge MA, USA; Department of Genetics, Harvard Medical School, Boston, MA, USA; Department of Neurology, Boston Children’s Hospital, Boston, MA, USA; Department of Biological Sciences, Purdue University, West Lafayette, IN, USA

## Abstract

Inflammation is increasingly recognized as important to neuropathology, including more classic neuroimmune disease as well as neurodegenerative and neuropsychiatric disorders. Interferons (IFN) are important mediators of central nervous system inflammation. Individuals appear to vary in susceptibility to neuroinflammatory pathology, suggesting that identifying human genetic modifiers of the neuronal IFN response might provide insight into disease pathophysiology. To identify potential modifiers, we stimulated neuronal “cellular villages” of iPSC-derived neurons from over one hundred donors with IFN-alpha (IFNa) or IFN-gamma (IFNg). We then correlated allele states of common variable SNPs to gene expression to identify hundreds of expression quantitative trait loci (eQTLs), many of which emerged specifically upon IFN treatment. We characterized the distinct but overlapping neuronal transcriptional responses to IFNa and IFNg, and identified specific response QTLs. Functional annotation of STAT1 binding to the genome in response to IFN stimulus identified STAT1 binding sites as enriched for response-regulating human genetic variation and also enabled identification of loci with IFN-dependent allele-specific binding of STAT1. These results demonstrate how human genetic variation can influence IFN-dependent mechanisms in neurons in disease-relevant ways.

## Introduction

Neuroinflammation plays a critical role in neurodegeneration, neuropsychiatric disorders and autoimmune encephalitis^1–3^. Interferons (IFNs), a key part of the inflammatory response, are important cytokines in different arms of the host defense and can contribute to central nervous system (CNS) injury. Uncontrolled high levels of interferon-alpha (IFNa) in diseases such as neuropsychiatric systemic lupus erythematosus (NPSLE) and Aicardi Goutières Syndrome (AGS), or of interferon-gamma (IFNg) in multiple sclerosis (MS), correlate with disease activity which can include seizure, neuronal cell death, brain atrophy, and severe disability^1,4,5^. Interferons have also been shown to have important functions in neural cells, such as IFNg’s effect on dendritic spine density^6^, neurite outgrowth^7^ and neuronal cell death^8–10^. Type I interferons such as IFNa have been shown to have multiple functions in neurodegeneration and neuropsychiatric disorders^3^ including promoting neuronal synaptic loss in Alzheimer’s models^11^ and also in neuronal survival^12^. Together, these relationships suggest interferon signaling in the brain contributes to neural physiology and pathology.

CNS neuroinflammatory diseases are also notable for significant variability in their disease course across individuals^13,14^. This variation is likely multifactorial, spanning both extrinsic factors (such as infectious exposures) and intrinsic (genetic) factors^13^. While prior work has uncovered genetic variability in the immune system in neurological disorders^15,16^, less is known about how human genetic modifiers affect how neurons and other brain cells respond to cytokines^17,18^. Elucidation of these modifiers could in principle reveal novel mechanisms of inflammation-mediated neuropathology and suggest new therapeutic avenues.

Significant human genetic variation has been identified at the level of single nucleotide polymorphisms (SNPs)^19^, which can lead to phenotypic diversity by modifying gene expression. Prior studies such as the Genotype-Tissue Expression (GTEx) project have identified thousands of common SNPs associated with RNA expression of nearby genes, relationships known as expression QTLs (eQTLs), that are increasingly recognized as having functional biologic outcomes^20–24^. However, these studies require DNA samples from thousands of donors and are usually limited by the ability to attribute a single gene in the associated region to pathobiology. Mechanistic validation is challenging, as is identifying the cell type and the context in which the gene is important. This has begun to be addressed in immune cell populations^16,25,26^, but has been limited in neural cells, which are harder to access^27^.

A significant limitation to assessing the effect of natural human genetic variation in response to a specific stimulus in neural cells has been the lack of a scalable system for analyzing genetically diverse sets of human cells under identical growth conditions. We turned to induced pluripotent stem cell (iPSC)-derived neurons as an experimental system uniquely suited to enable the manipulation and stimulation of human neurons derived from many donors^28,29^. To limit technical variation and to be able to correlate genotype variation with cellular phenotypes, we used the “cell village” approach^20,30,31^, in which we culture iPSC-derived neurons from many donors together. By placing all donors’ cells in a common culture environment, this approach minimizes technical sources of variation and maximizes power to recognize true genetic effects with modest numbers of individuals. We analyzed the response of iPSC-derived neurons (from a total of 102 distinct donors) to IFNa and IFNg and found interindividual variation in this response at multiple levels, including hundreds of sites of IFN-specific human genetic variation.

## Results

### Interindividual variation in the transcriptomic response to IFNs

In order to examine how individual donors’ neurons respond to IFNs and how these responses might vary, we generated iPSC-derived excitatory neurons from 102 previously genotyped human donors^32^ (Figure 1a; Demographics in Figure S1a; Table S1), and stimulated them with IFNg for 24 hours or IFNa for 7 hours at doses and time points adapted from prior work demonstrated to produce robust responses in other cell types^33,34^. After removing donors with insufficient representation (Methods), there were 80-88 unique donors per condition with 64 donors shared across all three conditions (Figure S1b; Table S1). We analyzed their single-cell RNA-seq profiles and used transcribed SNPs to assign each cell’s donor, then combined all cells from a given individual to generate a “metacell” to quantify average gene expression changes for each donor. We then examined overall induction of gene expression in response to IFN treatment in our system.

**Figure 1:**
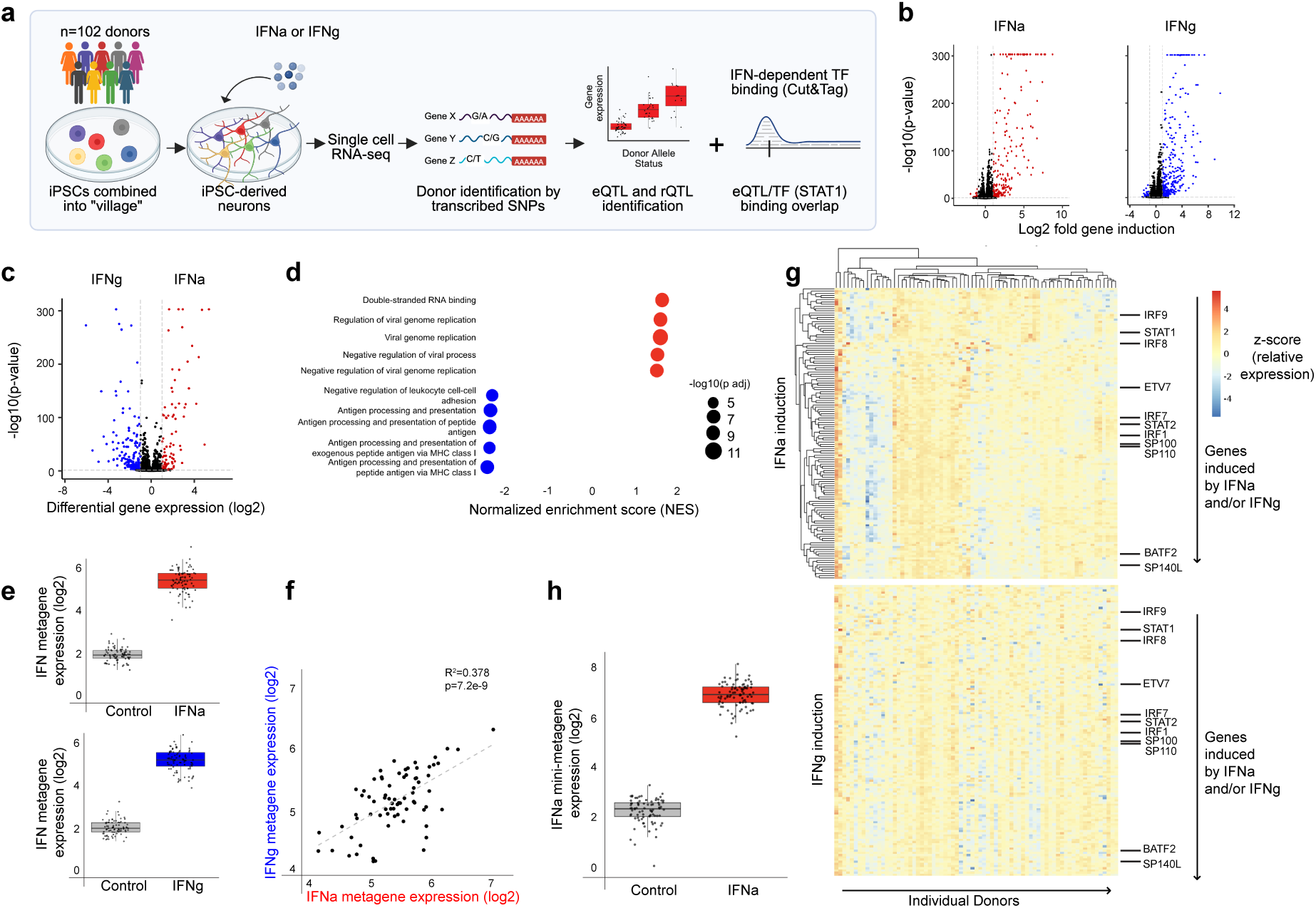
Interindividual variation in the transcriptomic response to IFNs. **a**) Experimental overview. iPSC-derived neurons from 102 donors (genotyped separately before analysis) were cultured together and treated with IFNa, IFNg or PBS (control). Cells were analyzed by single-cell RNA-seq, then assigned to the individual donors based on combinations of many transcribed SNPs. Later, in some analyses, genetic sequence variation and gene expression are correlated to identify eQTLs, and this is compared to transcription factor binding. **b**) Volcano plots show overall response to 7 hours of IFNa (left) and 24 hours of IFNg (right) treatment with robust gene upregulation. Data are from all donors, all cells (pseudo-bulked together). **c**) Differential gene-expression responses to IFNg and IFNa treatment. **d**) Gene set enrichment analysis comparing differentially expressed genes in IFNg versus IFNa treatment. Top five enriched by normalized enrichment score (NES) biological pathways shown (highest and lowest). **e**) IFN-response metagenes were generated by calculating mean expression of top induced genes with >4-fold induction (n=94 for IFNa, n=109 for IFNg). IFNa-response and IFNg-response metagene expression per donor in both control (gray) and IFN conditions is shown for IFNa (red) and IFNg (blue). **f**) IFN-response metagene expression for each donor in response to either IFNa (x-axis) or IFNg (y-axis). **g**) Relative expression (visualized as Z-score of the donor-specific measurements) of gene expression (log counts per million) of top induced genes (fold change >4) in response to IFNa (top) or IFNg (bottom). Each column represents a donor; each row represents a gene. Hierarchical clustering was performed for both donors and genes in the IFNa group, and this order was maintained to display IFNg data. **h**) “mini” metagene magnitude shown per donor in both control (gray) and IFNa-stimulated (red) conditions.

Interferons caused robust induction of gene expression changes, with 224 genes with IFNa and 354 genes with IFNg significantly induced more than 2 fold (FDR<0.05), and a smaller number of genes significantly decreased in expression (39 in IFNa and 69 in IFNg (Figure 1b; Table S2). IFNa and IFNg shared many similar pathways and target genes with 68 genes induced greater than 4 fold by both IFNs (Figure 1c; Table S2), but their responses also had distinct features (Figure 1c) including 26 genes induced greater than 4 fold in IFNa but not IFNg, and 41 genes induced greater than 4 fold in IFNg but not IFNa.

The unique gene induction profiles in response to either interferon family member were highly enriched for pathways (identified by Gene Set Enrichment Analysis) known to be important in the IFN responses of immune cells^35^ (Figure 1d), and also to have unique functions in neurons^7,12^.

The top pathways included “antigen processing and presentation of peptide antigen via MHC class I”, induced by IFNg and “double stranded RNA binding”, by IFNa.

To evaluate whether responses to IFN varied on a donor-by-donor basis, we first calculated the average expression in each donor of the most strongly induced genes (Table S3) as a “metagene” and compared their expression in control or IFN conditions. We found that though each donor’s cells responded robustly to IFNa or IFNg, the donors exhibited about 5 fold (for IFNa) to 10 fold (for IFNg) variation in expression levels of these IFN-induced genes (Figure 1e). Intriguingly, when we examined the per donor response, we found that overall donors whose cells responded most strongly to IFNg also tended to have the highest magnitude of average response to IFNa, suggesting underlying factors, potentially genetic, shape the magnitude of both responses (Figure 1f; R^2^=0.378, p=7.2e-9).

We next compared the response on a gene-by-gene level, to see if variability was driven by a few select genes, or overall differences in gene induction. Some donors more robustly induced IFN gene expression overall, where the relative expression of each gene correlated to the overall magnitude of response (Figure 1e, 1g). For donors with similar magnitude of responses to IFNa and IFNg, this seemed to be reflected across most genes instead of driven by a select few (Figure 1g). Thus, the correlation between IFNa and IFNg did not appear to be driven by one commonly induced gene in particular but by the differential expression of many target genes together.

IFN receptor expression level has been suggested to correlate with the magnitude of IFN response^36^. However, we did not find substantial correlation between the overall magnitude of the IFN response and IFN receptor expression at baseline (all R^2^<0.09). There was a subtle increase with IFN stimulation of the correlation of *IFNAR1* and in particular *IFNGR2* expression (which is itself induced by IFNg) (Figure S2a-b).

Additionally, subsets of donors appeared to share a stronger response in subsets of the IFN response that corresponded to specific modules of genes that were particularly strongly correlated in their responses (sets of adjacent rows in Figure 1g). Eleven genes encoding transcription factors (TFs) induced by IFNs were distributed throughout these modules (Figure 1g), nominating candidate regulators of these modules.

Finally, we considered clinical correlates of these responses, and in particular a 6-gene IFNa “mini-metagene” response that has been measured in patient blood as an interferon-stimulated gene signature in type I interferonopathies^37^. We found that this IFN response was also robustly induced in donor neurons, with 7.5-fold variation across individuals (Figure 1h; Table S3). This response also had minimal correlation with the IFNAR expression levels particularly at baseline (Figure S2c; *IFNAR1* R^2^=0.033, p=0.13; *IFNAR2* R^2^=0, p=0.98).

### Genetic determinants of IFN response and cis-eQTL discovery

We next leveraged the whole genome sequencing data from each donor to define IFN-responsive cis-expression quantitative trait loci (eQTLs). We identified hundreds of eQTLs, including 516 that reached statistical significance only with IFNa and/or IFNg stimulations but not in baseline conditions, and which we thus considered to be “IFN-responsive” (Figure 2a-c; Table S4). To further validate these eQTLs, we profiled gene expression in a second independently differentiated village of 36 donors (Table S1) and used a sign test to compare eQTLs across the two villages. We found that 95% of eQTLs identified from the first village had the same direction of effect (sign) of the associated allele in the two different villages. Intriguingly, some genes, such as *CASP7*, were robustly induced in both IFN conditions, but in an allele-dependent context in only one of the conditions (e.g. for *CASP7*, IFNg) (Figure 2c).

**Figure 2:**
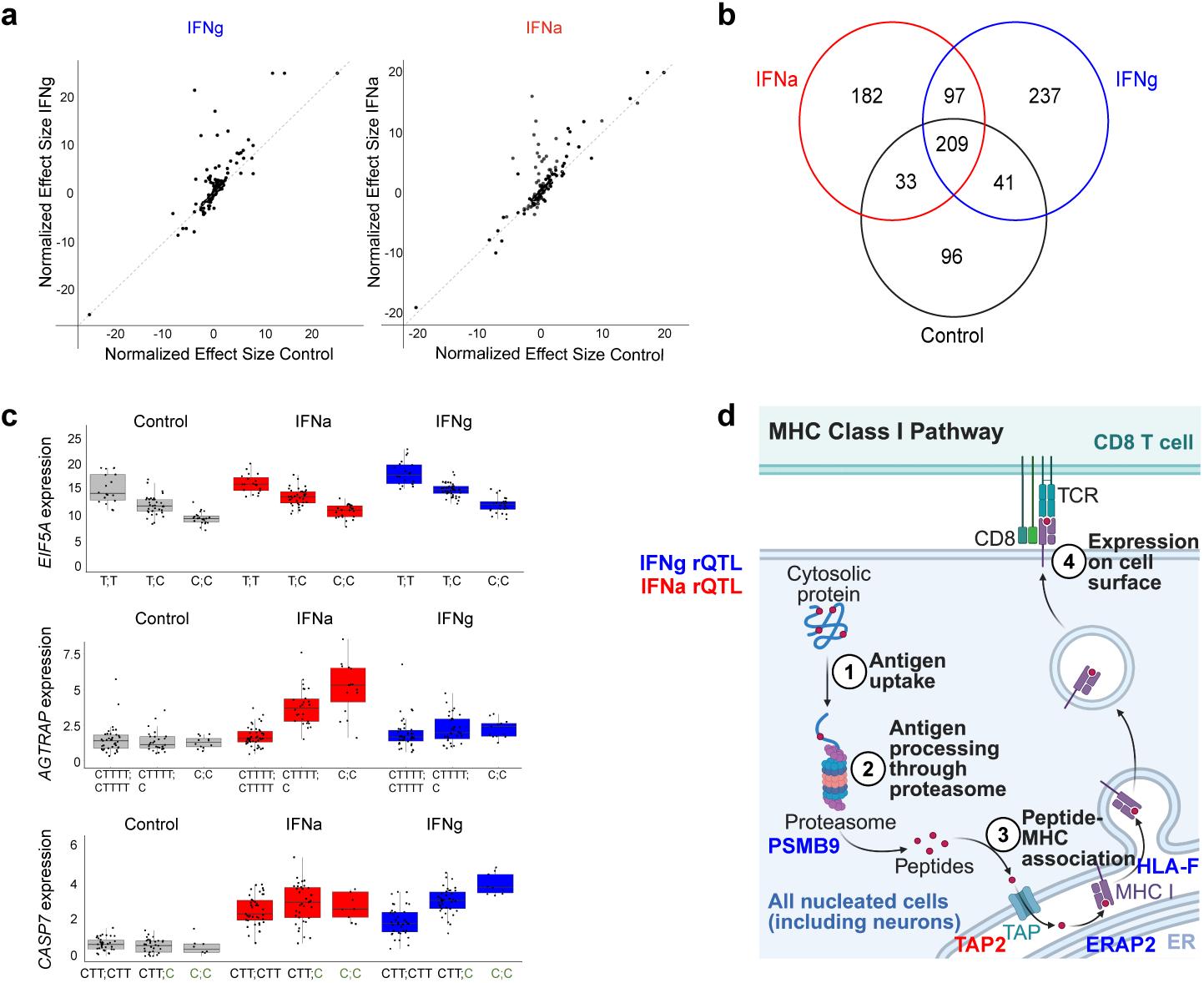
Genetic determinants of IFN response and cis-eQTL discovery. **a**) Comparison of normalized eQTL effect sizes in IFN versus control. Significant eQTLs in either IFN conditions are displayed, comparing their effect size normalized to respective gene expression in either control or IFN conditions. Off-diagonal effects are response-specific. **b**) All eQTLs and their corresponding genes with q-value <0.05 were considered, and diagram displays overlap in genes with eQTLs in each condition. **c**) Example eQTLs including those considered shared (top) and IFN-responsive (middle, bottom) **d**). Schematic of significant rQTLs found in IFNa (red) and IFNg (blue) response and their area of function in the classical MHC class I peptide presentation pathway.

We further defined response QTLs (rQTLs), which specifically examine the interaction between two stimuli and a specific SNP^38^. Using this more stringent approach, we found 5 rQTLs in the IFNaxControl (Table S5) and 11 rQTLs in the IFNgxControl conditions (Table S6). We note that the IFNg rQTLs were at loci encoding proteins expected to interact with each other significantly more than expected (stringDB^39^ protein protein interaction p-value 5.7e-5). Overall, IFN rQTLs were enriched in the MHC Class I presentation pathway (reactome FDR 0.0483) (Figure 2d; Table S5-S6) across multiple loci (*TAP2, PSMB9, HLA-F* on chromosome 6; *ERAP2* on chromosome 5).

### STAT1 binding underlies much interindividual variation in response to interferons

The vast majority of the eQTLs we identified involved SNPs in non-coding genomic segments with unclear function (Figure 3a). To functionally annotate these, we examined their overlap with transcription factor binding. While IFNa and IFNg use different primary transcriptional mechanisms, both employ Signal Transducer and Activator of Transcription 1 (STAT1) to regulate downstream gene expression^40^. Downstream of IFNa, STAT1 is preferentially engaged as part of the ISGF3 heterotrimer, which binds to interferon-stimulated response elements (ISRE)^40^. In IFNg-dependent signaling, STAT1 binds as a homodimer to its motif, the gamma interferon activation site (GAS) motif. Therefore, we sought to better characterize STAT1 genome-wide binding, and performed CUT&Tag for STAT1 in neurons under control conditions and after IFNa and IFNg stimulation. IFN stimulation elicited robust increases in STAT1 binding (Figure 3b). We identified high-confidence areas of STAT1 binding across the genome in neurons, with 1175 genomic regions (peaks) bound by STAT1 in IFNa stimulus conditions and 7180 bound by STAT1 in IFNg conditions (compared to just 377 sites in the control condition). We further found that the GAS/STAT1 DNA binding motif (RTTTCCNGGAAA) was the most enriched DNA-sequence motif in STAT1 binding sites in response to IFNg (p<1e-1396, motif rank 1), and that the ISRE (GAAACCGAAA) was one of the top motifs in the IFNa STAT1 binding sites (p<1e-36, motif rank 7), consistent with the binding mechanisms known for this transcription factor (Figure 3c)^40^.

**Figure 3:**
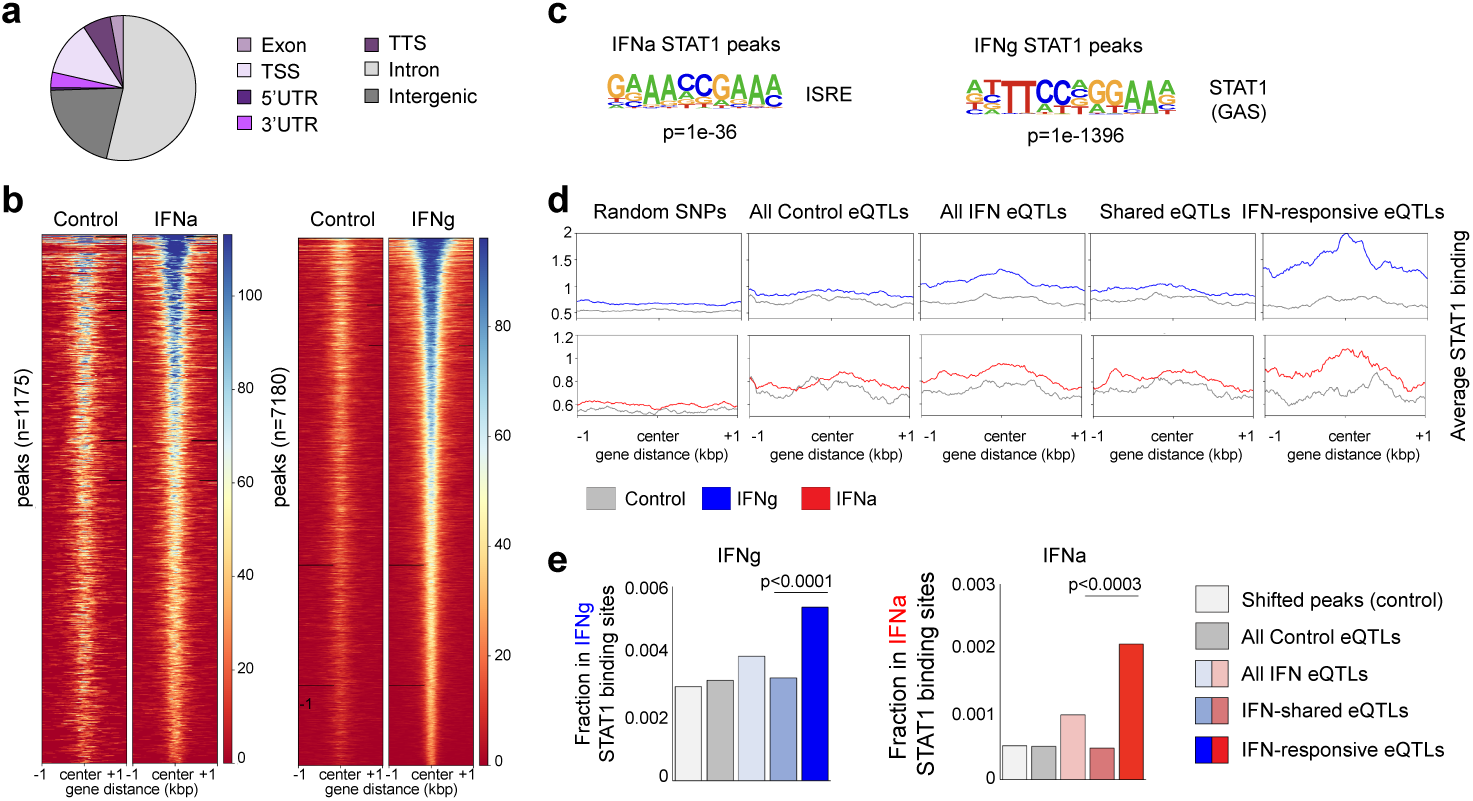
STAT1 binding underlies much interindividual variation in response to interferons. **a**) Distribution of all eQTLs found (in any condition) in the genome. **b**) STAT1 binding across the genome in response to either IFNg (right) or IFNa (left). **c**) Enriched motifs in STAT1 peaks relevant to STAT binding. **d**) average mean read density of STAT1 binding (measured using Cut&Tag) at SNPs and eQTLs identified in control and IFN-response conditions. **e**) Overlap of eQTL categories with STAT1 binding sites. Chi-squared test performed on IFN-shared eQTLs and IFN-responsive eQTLs.

We next asked whether IFN-responsive eQTLs were more likely to be at sites of STAT1 binding (Figure 3d, 3e). First, we evaluated the mean STAT1 binding signal in the presence of IFNg (Figure 3d, top) or IFNa (Figure 3d, bottom) as compared to control. At baseline, there was minimal induction of STAT1 binding in randomly selected SNPs in the genome, or at eQTLs identified in the control condition. Among eQTL SNPs identified in the IFN-treated condition, there was an appreciable increase of STAT1 binding (Figure 3d). These eQTLs in principle include a mixture of (i) eQTLs shared with the control condition, and (ii) eQTLs specific to the response condition, “IFN-responsive”. When we separated the eQTLs into these two groups, we observed that the increase in IFN-dependent binding of STAT1 arose entirely from the IFN-responsive eQTLs (Figure 3d).

As an alternative analytical approach, we estimated the fraction of eQTLs that fell in a STAT1 binding site (Figure 3e). IFN-responsive eQTLs were significantly more likely to overlap with STAT1 binding sites than IFN eQTLs that were shared with the control condition. The enrichment of these eQTLs at STAT1 binding sites suggests that areas of STAT1 binding and STAT1 signaling pathways are an important source of human genetic variation in the neuronal interferon response.

### Allele-specific and interferon-dependent function at a CASP7 eQTL

The strong enrichment of STAT1 binding sites among IFN-responsive eQTLs (Figure 3) allowed for the selection of eQTLs in functionally relevant binding sites to examine how they may affect transcriptional mechanisms. One of these was at the *CASP7* promoter, rs10553596 (Figure 4a), and was found to be an IFNgxPBS rQTL. Rs10553596 is a common variant with a major allele characterized by the presence of CTT and a minor allele with just C. The minor allele has been found to be protective against the development of Alzheimer’s disease^41^.

**Figure 4:**
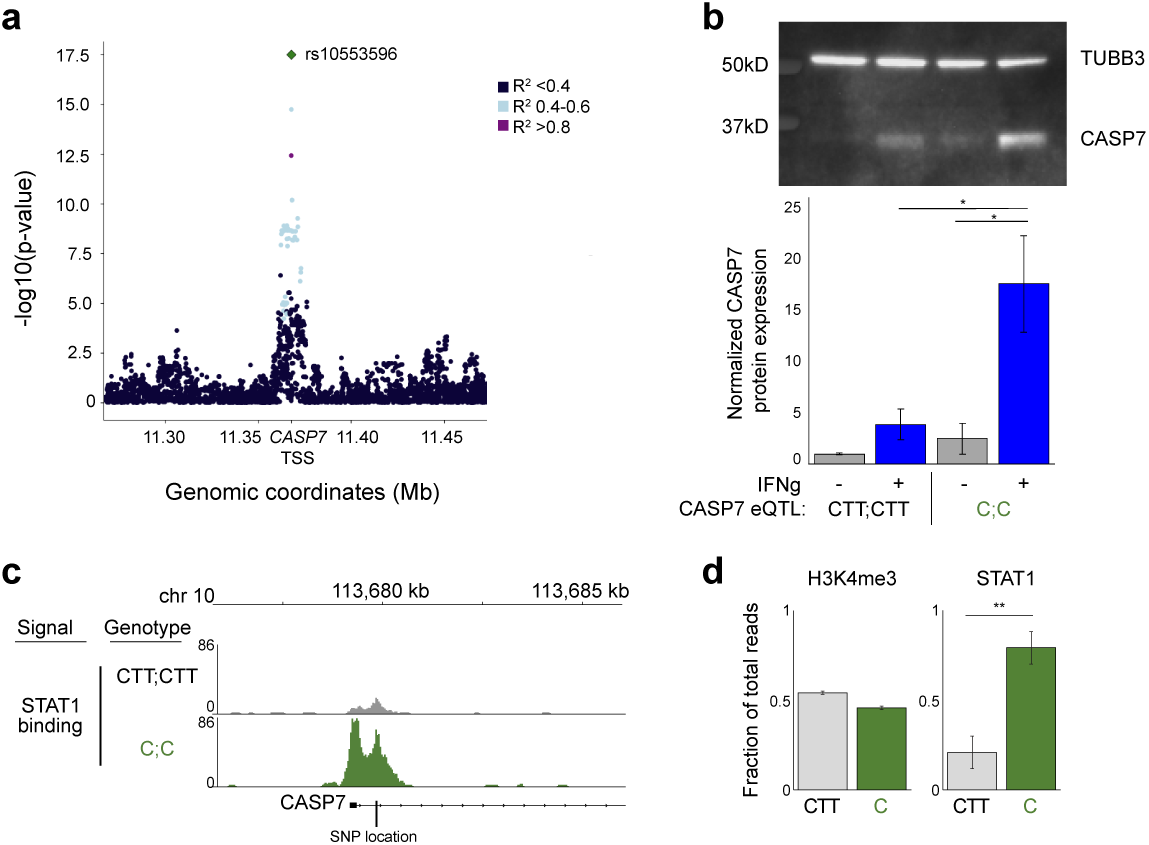
Allele-specific and interferon-dependent function at a *CASP7* eQTL. **a**) SNPs within 500kb of the lead *CASP7* eQTL are shown colored by R^2^ value with respect to the lead *CASP7* eQTL (green). **b**) Top, example western blot for CASP7 with TUBB3 as loading control. Bottom, protein quantification normalized to TUBB3 expression (n=3 independent pairs) *p<0.01 by student’s t-test. **c**) STAT1 binding at the *CASP7* promoter in cells carrying two copies of the major (CTT;CTT) or minor (C;C) allele. Signal is normalized to total number of reads. **d**) Relative STAT1 Cut&Tag read density across CASP7 eQTL in cells heterozygous at the allele (CTT; C) (n=4). **p<0.0001 by chi-squared test (for each experiment). H3K4me3 read density across the same site as STAT1 binding, shown as control (n=2; p-value for each chi-squared test = 0.04 and 0.52).

We first asked whether the effect of this common variant manifested on a protein level as well. Neurons with the C allele indeed induced the CASP7 protein more strongly (in response to IFNg) than neurons with the CTT allele did (Figure 4b). The minor allele (C) is computationally predicted to provide a better STAT1 binding site^42^. Furthermore, STAT1 is primarily an activating transcription factor^40^, and increased STAT1 binding at the minor allele (for which we observed higher RNA and protein expression) would be consistent with this. To test this, we performed Cut&Tag in neurons from donors homozygous for either the major allele (CTT) (Figure 4c, top) or minor allele (C) (Figure 4c, bottom). We identified a peak corresponding to STAT1 binding overlying this eQTL, and confirmed that donors homozygous for the minor allele had notably more STAT1 binding (Figure 4c).

To further test the idea that STAT1 binds to this site in an allele-specific manner, we examined STAT1 binding in donors heterozygous at this allele, and again found significantly increased binding at the minor allele within the same cells (Figure 4d). As a control, we looked at H3K4me3 binding at this variant and did not find a similar skew, nor did we find consistent significant differences in allele binding. Overall, these data demonstrated that the minor allele resulted in more STAT1 binding and *CASP7* mRNA and protein production.

## Discussion

In this work, we examined how human neurons from a total of 102 donors varied in their response to interferons, to explore why individuals may present with different neuroinflammatory phenotypes. We found variation in the response at multiple levels, including in the overall magnitude of the response as well as among subsets of genes, suggestive of nodes of regulation that may correspond to different arms (modules) of the IFN response. One intriguing possibility to explore in the future is that variation in transcription factor induction and function could serve as these nodes of interindividual modular variability. Two distinct modules of TFs induced by type I interferons were recently identified in T cells, one characterized by STAT/IRF family members and the second by BATF/AP1 family members^43^. We identified similar transcription factors in the response to IFNa (STATs, IRFs, BATF2) suggesting neurons may employ similar approaches.

The discovery of eQTLs and rQTLs in the neuronal response to IFNs, and the identification of neuron-wide STAT1 binding, provides the opportunity to functionally annotate the results of genome-wide association studies. We focused on common variants with a minor allele frequency of greater than 5%, thus relevant for large fractions of the population. As an example, we highlighted a locus at the *CASP7* promoter associated with Alzheimer’s risk^41^, rs10553596, and showed that it has allele-specific and interferon-dependent function. The frequency of the minor allele at rs10553596 globally is 0.29^44^. Therefore, 8.4% of the global population is homozygous for this minor allele. *CASP7* encodes executioner caspase 7, important in the final stages of apoptosis^45^. Notably treatment with IFNg was previously found to increase neuronal susceptibility to Amyloidβ(1-42) induced cell death^10^. A reduced ability of cells to complete apoptosis can lead to stressed cells choosing alternative pathways to cell death such as necroptosis^46^, an inflammation-generating pathway that has been found to be increased in brains from those with Alzheimer’s disease^47^. Intriguingly, higher serum IFNg has also been recently associated with slower cognitive decline in elderly patients^48^. Taken together, this might explain a potential protective role for IFNg-induced increased *CASP7* expression in Alzheimer’s neuropathology, where having two copies of the minor allele could in principle increase the propensity of neurons to undergo apoptosis and thus decrease the propagation of inflammation.

Some of these eQTLs, including *CASP7*, have been identified in monocytes in response to IFNg, although in this cell type this is not an IFN-specific eQTL^25^. However, the robust, IFN-dependent allele-dependent expression in neurons of these eQTLs opens up the possibility that they may not be immune cell-specific, but may modify function across cell types in different contexts.

Antigen processing and presentation on class I MHC may be another pathway that has context-dependent specificity in neurons, as class I MHC has been shown to be important in both synapse development^49^ and neurite outgrowth^7^, and neurons expressing MHC class I could activate CD8+ T cells which have been implicated in autoimmune encephalitis^50–52^. Class I MHC is expressed on human neurons in inflammatory contexts^7^. While class I MHC haplotypes themselves (*HLA-ABC*) are known to be an important risk factor for autoimmunity including neuroinflammatory disease^53,54^, variation within the pathway that contributes to differential self-peptide generation or loading could further modulate disease risk. Another IFNg rQTL, *BTN3A2*, has been implicated in gamma-delta T cell activation^55^, again suggesting specific variability in the ability to activate T cells as an important source of interindividual variation in the risk for T cell-mediated neuroinflammation.

### Limitations and Future Directions

The specificity of stimulus and cell type in this system, together with minimizing technical variability, allows for the identification of hundreds of eQTLs as well as rQTLs from just 102 donors. However, this may limit the generalizability of these findings, and further work examining different contexts, stimuli and time points will help clarify where stimuli-specific eQTLs and rQTLs are important. Furthermore, complementary work to examine the cell biological implications of these eQTLs and rQTLs will be important. However, this initial evaluation highlights how these sites of human genetic variation may be important in neuronal biology and provide an additional layer of modulation of neuroinflammatory disease risk.

## Contributions

M.M.A designed the study, analyzed the data, performed experiments and wrote the manuscript with input from other authors. S.B. N.P and D.M. analyzed the data. M.T. generated single cell RNA sequencing data for analysis. A.J. and I.M. prepared neuron cultures with assistance from D.H. C.J.M. and C.T.Y. assisted with cut and tag development and sequencing. G.G. calculated linkage disequilibrium for eQTLs. A.M., K.I. and M.H. contributed to project management and sequencing. S.M. and R.N. helped design and supervise the study and contributed to revisions of the manuscript.

These authors contributed equally: Sonia Bolshakova, Noah Pettinari.

## Conflicts of interest

The authors declare no competing interests.

## Acknowledgments

This work was supported by NIH grants R25NS070682 (M.M.A.), 5K12NS098482 (M.M.A.), U01MH115727 (R.N., S.A.M.), Boston Children’s Hospital Office of Faculty Development Grant (M.M.A.), the Broad Institute Next Generation Award (R.N.) and the Stanley Center for Psychiatric Research Gift (R.N.). BioRender was used in the generation of Figure 1a and 2d.

## Data Availability Statement

Data is available at ANVIL. Users should visit https://explore.anvilproject.org/datasets/db7caf73-bb61-4f40-bbf1-fa1489f71068 and click “Request Access”.

## Supplementary information

*Supplemental Figures and Tables*

Figures S1-S2 (both related to Figure 1).

Table S1 (Related to Figures 1-4): Metadata for all iPSC lines used and their use per experiment as well as *CASP7* allele status for relevant lines.

Table S2 (Related to Figure 1b-d): Differential gene expression in control, IFNa and IFNg conditions across all donors.

Table S3 (related to Figure 1e-h): Genes included in metagenes for IFNa or IFNg and their correlation with their respective metagene expression.

Table S4 (Related to Figures 2-3): Summary statistics for eQTL analysis in control, IFNa or IFNg conditions.

Tables S5-S6 (both related to Figure 2d): Significant identified rQTLs.

## Methods

### Cell lines

Human iPSCs were obtained from multiple sources. A list of all cell lines used in this study and which experiments they were used for is listed in Table S1. The majority of cell lines are available from the California Institute of Regenerative Medicine (CIRM) or the NIMH Repository and Genomics Resource (NRGR). Some Cut&Tag assays were performed using a WTC11 line expressing inducible NGN2 previously described^56^ (Gift of Michael Ward, commercially available now at Coriell (GM29371)).

### Cell culture

iPSCs were grown and differentiated into neurons adapted from a previously described protocol^32,57^. Briefly, iPSCs were differentiated by doxycycline-mediated induction of NGN2 and small molecules, and grown as of induction day 4 in Neurobasal Media (Thermofisher) with CNTF, GDNF and BDNF (Stemcell Technologies). Experiments were performed on Day 28-30 of induction. For RNA experiments neurons were grown on pre-plated mouse glia. For protein experiments and Cut&Tag, neurons were grown as a monoculture. Cells were either stimulated with PBS (control), IFNa (50ng/ml, 7 hours, Sigma #H6166) or IFNg (50ng/ml, 24 hours, Sigma #I17001) unless specified differently below.

### Genotyping iPSC lines

Whole-genome sequencing (WGS) data was generated for each donor used in the experiment via Illumina PCR-Free Human WGS - 30x v2, via the Broad Institute’s Genomics Platform. After sequencing, the raw cram files were joint-called using the GATK (v.4.5.0) WGS variant calling workflow.

Concordance for each donor was validated using the KING relationship inference software (v2.3.0) against ground truth genotype data obtained from the stem cell sources (GSA for CIRM lines obtained from California Institute of Regenerative Medicine/CIRM and WGS for McLean-Levy lines obtained from Columbia University) to identify putative sample swaps, duplicates, or mixed samples between cell lines from different donors. All sequenced cell lines were confirmed concordant with their donor of origin.

After validating donor concordance, the VCF was prepared according to Dropulation standards^20^.

### Cell village creation, scRNA-sequencing and donor assignment

IPSCs from each donor were mixed in equal proportions (1M cells per donor) and differentiated as a pool following the excitatory neuron differentiation method described earlier. At D30 of the differentiation, following IFN stimulation, neuron villages were harvested, and 60,000 cells were prepared using 10X Chromium Single Cell 3’ Reagents v3 and sequenced on an Illumina NovaSeq 6000 with an S2 flow cell, generating paired-end reads of 2 x 100 bp. Raw sequencing data were aligned and processed following the Drop-seq workflow^58^. Human reads were aligned to the GRCh38 reference genome and filtered for high-quality mapped reads (mapping quality ≥ 10). To determine the donor identity of each droplet, variants were filtered through multiple quality controls, ensuring only high-confidence A/T or G/C sites were included in the VCF files. Once the single-cell libraries and VCF reference files were filtered and quality-checked, the Dropulation algorithm (v.2.5.3) was applied. This algorithm analyzes each droplet (or cell) independently, assigning a probability score to each variant site based on the observed versus expected allele. Donor identity is determined by computing the diploid likelihood at each UMI, summed across all sites, to identify the most likely donor for each droplet. To ensure consistency in differential expression testing downstream, we remove any cells which are clustered separately by individual donors. After donor assignment, digital gene expression (DGE) matrices from each sequencing library were merged into a single DGE matrix for downstream analysis. Donors whose expression was less than 0.3% of the population were removed from downstream analyses as per Dropulation workflow.

### Differential expression analysis

For differential gene expression analysis (DGEA) on the village data, expression data was pseudobulked by donor by summing the counts per cell for each gene within each donor. The resulting pseudobulked matrix was then used for the downstream differential expression analysis. Differential expression analysis of pseudobulked scRNA-seq data was performed on human neurons using a combination of the DESeq2 (v1.34.0) and limma (v3.48.3) packages in R (v4.1). Read counts were first normalized for library size using DESeq2’s internal size factor normalization and voom was used to produce log2-transformed normalized gene expression estimates. Genes with low read counts (<= 10 reads in at least one sample and total reads <= 15) were removed using standard parameters (described above) with the filterByExpr function in edgeR (v3.36.0) prior to library normalization. For differential expression analysis of the bulk effect of interferon stimulation independent of genetic background, a single factor model was used with treatment as the covariate (∼0 + treatment). P-values for differential expression were adjusted using an FDR of 5% and Bonferroni-Hochberg multiple testing correction. Gene set enrichment analysis was performed on the differential expression analysis results from limma-voom using the fgsea package (v1.18.0) in R using 10,000 permutations and the t-statistics as the ranking metric. The C5 Gene Ontology collection (v7.2) from the Molecular Signatures Database was merged with the SynGO (release 20210225) biological process (BP) and cell component (CC) gene lists and was used with a custom gene set comprising genes implicated in schizophrenia genetic studies of humans, including both genes at 1-2 gene loci from the SCZ GWAS (PGC3) and genes containing rare coding variants (SCHEMA; FDR<0.05).

Enriched gene sets were then adjusted for multiple testing using Bonferroni-Hochberg correction at an FDR of 5%.

### Metagene analysis

For the metagene analysis, we aggregated gene expression counts for two different lists of genes: a “small” metagene for IFNa, and a “logFC” metagene for both IFNa and IFNg, which are outlined in Table S3. Metagene values for each donor were computed from the pseudobulked DGE as the mean logCPM for all genes within each metagene. Metagene values were compared both between metagenes and between each gene comprising each metagene. 100/109 genes in the IFNg metagene and 91/94 genes in the IFNa metagene had a significant correlation (p<0.05) with the overall metagene (Table S3).

### QTL analysis

General eQTL analysis was performed using Matrix eQTL^59^. Prior to running eQTL analysis, pseudobulked counts were normalized such that genes whose expression fell in the bottom 50% of genes expressed were removed, and each donor had a total of 100,000 counts. Additionally, we included covariates for each run of Matrix eQTL detailing the following covariates for each donor: sex, age at iPSC collection, and the first 3 genetic PCs (computed from a PCA on the input VCF file). For donors whose age was unknown at the time of iPSC collection, their age was set to the median age of their reported sex. Additionally, for cis-eQTL gene expression analyses, we also included 10 latent factors to capture gene expression batch effects, calculated from PEER (v1.0). All eQTL analysis on the larger village was performed using a minor allele frequency (MAF) of 5%.

We performed cis-eQTL analysis for each treatment (IFNa, IFNg, and PBS) at windows of 10kb. Because our smaller village (n=35 for PBS, 36 for IFNa, IFNg) was less than half the size of our larger village, we ran cis-eQTL analysis at a MAF of 20% due to limited statistical power for eQTL detection. Upon repeat with the main village, MAF of 5% was used. Permutation testing was conducted using 10,000 permutations. Permuted p-values were corrected using Bonferroni-Hochberg multiple-testing correction at an FDR of 5%.

Sign tests were run to validate eQTL concordance by comparing the direction of effect sizes between our two villages. There were 299 overlapping eQTLs in IFNa, 358 in IFNg and 225 in PBS. A positive sign test was if the direction of effect was consistent. In each treatment, we obtained a concordance rate >90%, adding support that batch effects were not impeding our ability to detect eQTLs.

For response QTL (rQTL) discovery, we leveraged the interaction mode of the tensorQTL (v1.0.9) package in Python^38^. The same covariates were used as for Matrix eQTL analyses as well as the same 10kb windows. Expression, covariate, and genotype data for treatment conditions (IFNa and IFNg) and control (PBS) were concatenated and labeled accordingly. rQTLs were nominated based on significant (p<0.05) genotype-treatment interaction p-values. Significant rQTLs were then determined if the Benjamini-Hochberg adjusted p-value was below 0.05.

eQTL localization in the genome was performed by first combining eQTLs across conditions, and removing duplicate eGenes for a total of n=895. These were then annotated using the annotatepeaks.pl function in Homer^60^ with the -genomeOntology option and hg38.

Linkage disequilibrium estimates were computed using PLINK^61^ from 1000 Genomes project high coverage^62^ genotype data for 503 samples labeled as European. Computation was run using PLINK options “--ld-snp <VARIANT_ID> --r2 dprime with-freqs --ld-window 100000 --ld-window-kb 1000”.

### Western blotting

Neurons were stimulated with IFNg (50ng/ml) for 48 hours and then lysed in RIPA with protease inhibitor (Roche), then processed into 4X Sample buffer with Dithiothreitol as a reducing agent. Samples were run on NuPage Bis-Tris 4-12% gels (Thermofisher), transferred using iBlot2 (Thermofisher), and membranes were probed with anti-TUBB3 antibody (1:1000, Cell Signaling Technologies, #5568) or anti-CASP7 antibody (1:1000, Abcam, #AB32522). Secondary antibody rabbit anti-HRP (1:10000) was used for chemiluminescence based readout. ImageJ was used for quantification.

### Cut&Tag

Cut&Tag^63^ was performed using Epicypher kit 2.0 with either 0.5μg anti-STAT1 (ProteinTech, #10144-2-AP) or 1μg H3K4me3 (Active Motif, #39916) with IgG control antibody from the kit. 100k nuclei were used per reaction. DNA input was normalized per sample before multiplexed sequencing. Sequencing was performed using the Nextseq550 mid-output kit with 150 cycles (Illumina, #20024904). 4-10M paired reads were obtained per sample used for peak calling. Overall 4-16M reads were obtained per sample.

### Cut&Tag computational analysis

Reads were mapped using Samtools^64^ to hg38. Peak calling was performed using the callpeaks function in Macs2^65^ with genome size specified as human. “High confidence” IFNa and IFNg STAT1 peaks were defined as those that were present in two separate bioreplicates, but not overlapping with IgG control antibody peaks.

For eQTL and peak overlaps, eQTL cohorts were defined as those eQTLs with a q-value <0.05. In order to account for uncertainty in lead SNP determination, all SNPs with a p-value within two orders of magnitude of the lead eQTL SNP were also included in the overlap analysis SNPs in Figures 3c and 3d, provided their p-value was of at least nominal significance (p<1e-5). Random SNPs were chosen in equal number to the number of “All IFN eQTLs” SNPs used in experimental overlap analysis. These cohorts of SNPs were also assessed for overlap with high confidence STAT1 peaks using BEDTools^66^.

### Peak Visualization

Data was processed using the computeMatrix.pl function of deepTools^67^. In each case the genomic locations provided (either high-confidence peaks, or SNPs in described categories) were set as the center of the data with a window of +/− 1kb. Heatmaps or average expression around the SNP categories are displayed for one bioreplicate for control and IFN conditions. Heatmap was generated using plotHeatmap function with deepTools^67^ sorting the peaks based on the IFN condition signal. Otherwise default settings used. Profile of IFN binding at described SNP categories was generated using plotProfile.pl in deepTools also using otherwise default conditions.

### Motif Enrichment

Motif enrichment in high confidence STAT1 peaks was obtained using the findMotifsGenome.pl function of Homer^59^ with the following parameters: -size given -S 20 -noknown.

## Notes

### Competing Interest Statement

The authors have declared no competing interest.

