## Supplementary material for "Human genetic variation shapes the response of neurons to interferons": Figure S1 and S2

### Figure S1: Characterization of donor iPSCs used in experiments.

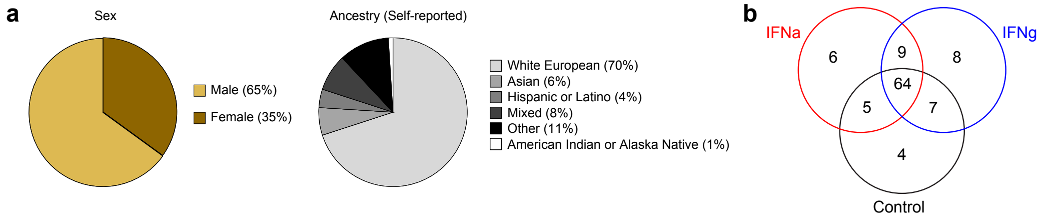


**a)** Demographics of donors. **b)** Overlap of donors across conditions. n=102 total donors across three conditions.

**Figure S2: Correlation of IFN receptor expression level with magnitude of IFN response.**

**a**

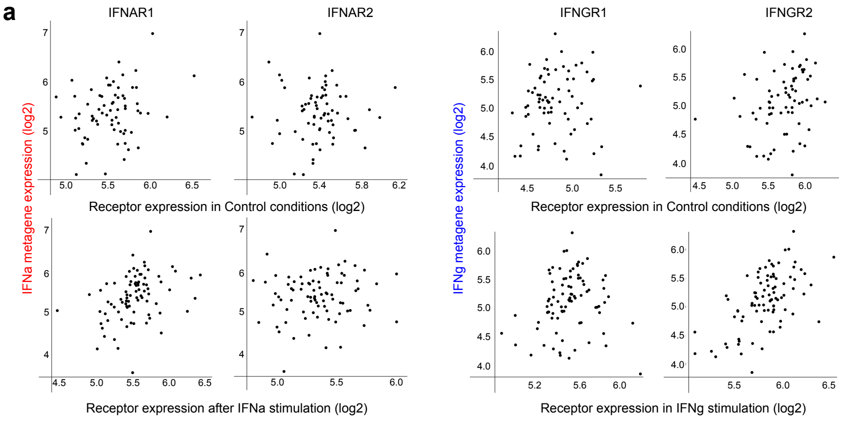


**b**

Coefficient of determination ($R^2$)

| | IFNAR1 | IFNAR2 | IFNGR1 | IFNGR2 |
|---|---|---|---|---|
| Control | 0.067 | 0 | 0.006 | 0.089 |
| IFN | 0.126 | 0.009 | 0.011 | 0.348 |

Most correlated (1) — $R^2$ — Least correlated (0)

Significance of correlation (p-value)

| | IFNAR1 | IFNAR2 | IFNGR1 | IFNGR2 |
|---|---|---|---|---|
| Control | 0.032 | 0.93 | 0.51 | 0.011 |
| IFN | 0.00094 | 0.38 | 0.34 | 1.50E-09 |

Most significant (0) — p-value — Least significant (1)

**c**

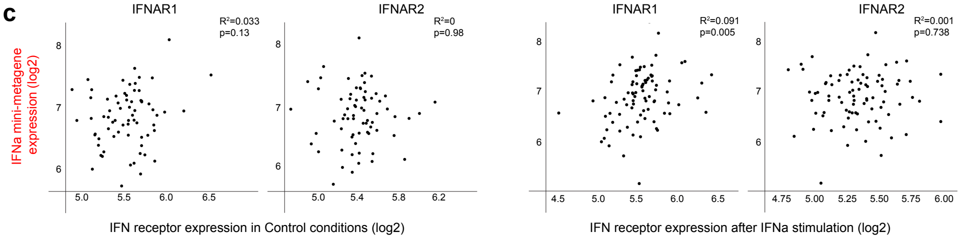


**a**) Dot plots demonstrating correlation of each IFN metagene with its respective IFN receptor (both subunits) on a donor-by-donor basis for expression of the IFN receptors in either control or respective IFN conditions. **b**) Summary of coefficient of determination (based on Pearson's correlation) and associated p-value for each metagene-receptor combination tested. **c**) Correlation of IFNa "mini" metagene expression after IFNa stimulation with RNA expression of *IFNAR1* and *IFNAR2* in either baseline or IFNa stimulation conditions. Coefficient of determination and p-value for association demonstrated for each combination.
