## Supplementary material for "Human genetic variation shapes the response of neurons to interferons": Table S5 and S6

**Table S5: Significant rQTLs found comparing the IFNa and Control condition (IFNaxControl).**

| Gene | variant_id | adjusted p-value |
|---|---|---|
| *AGTRAP* | chr1:11731850:CTTTT:C | 9.87E-12 |
| *TAP2* | chr6:32838799:C:T | 0.0104571 |
| *C8orf59* | chr8:85209076:G:T | 0.0307335 |
| *RPS26* | chr12:56042145:C:G | 0.00505825 |
| *SLFN5* | chr17:35244527:G:A | 1.04E-08 |

**Table S6: Significant rQTLs found comparing the IFNg and Control condition (IFNgxControl).**

| Gene | variant_id | adjusted p-value |
|---|---|---|
| *PDIA6* | chr2:10796583:T:A | 0.00665401 |
| *LAP3* | chr4:17598271:G:T | 5.72E-06 |
| *AC006160.1* | chr4:17579332:G:A | 0.0303002 |
| *ERAP2* | chr5:96916885:T:C | 2.08E-10 |
| *BTN3A2* | chr6:26358052:A:AAATAATAATAATAATAAT | 0.000406467 |
| *HLA-F* | chr6:29723313:T:A | 0.00360495 |
| *PSMB9* | chr6:32854082:C:A | 0.00371592 |
| *C8orf59* | chr8:85209076:G:T | 0.00675732 |
| *CASP7* | chr10:113679881:CTT:C | 5.74E-13 |
| *PRPH* | chr12:49291306:T:C | 0.00024032 |
| *MMP25-AS1* | chr16:3057948:C:T | 0.00943921 |
